# Histone acetyltransferase inhibition rescues differentiation of emerin-deficient myogenic progenitors

**DOI:** 10.1101/437343

**Authors:** Katherine A. Bossone, Joseph A. Ellis, James M. Holaska

**Author notes:** Correspondence, **Corresponding Author:** James M. Holaska, Department of Biomedical Sciences, Cooper Medical School of Rowan University, MEB 534, 401 South Broadway, Camden, NJ 08103. **Author contributions:** Conceptualization, methodology, and validation: J.M.H.; Formal analysis: J.M.H. and K.A.B.; Investigation: K.A.B. and J.A.E.; Resources: J.M.H.; Data curation: J.M.H., K.A.B., and J.A.E.; Writing-original draft: J.M.H.; Writing-reviewing and editing: J.M.H., K.A.B., and J.A.E.; Visualization: K.A.B. and J.M.H.; Supervision: J.M.H.; Project administration: J.M.H., K.A.B., and J.A.E. Funding acquisition: J.M.H. **Ethical Publication Statement:** We confirm that we have read the Journal’s position on issues involved in ethical publication and affirm that this report is consistent with those guidelines.

## Abstract

**Introduction:** Emery-Dreifuss Muscular Dystrophy (EDMD) is a disease characterized by skeletal muscle wasting, major tendon contractures, and cardiac conduction defects. Mutations in the gene encoding emerin cause EDMD1. Our previous studies suggested emerin activation of Histone Deacetylase 3 (HDAC3) to reduce Histone 4-Lysine 5 (H4K5) acetylation (ac) is important for myogenic differentiation.

**Methods:** Pharmacological inhibitors (Nu9056, L002) of histone acetyltransferases targeting acetylated H4K5 were used to test if increased acetylated H4K5 was responsible for the impaired differentiation seen in emerin deficient myogenic progenitors.

**Results:** Nu9056 and L002 rescued impaired differentiation in emerin deficiency. SRT1720, which inhibits the NAD^+^-dependent deacetylase Sirtuin 1 (SIRT1), failed to rescue myotube formation.

**Discussion:** We conclude emerin regulation of HDAC3 activity to affect H4K5 acetylation dynamics is important for myogenic differentiation. Targeting H4K5ac dynamics represents a new strategy for ameliorating the skeletal muscle wasting seen in EDMD1.

## INTRODUCTION

The nuclear envelope is composed of two lipid bilayers, the outer nuclear membrane, which is contiguous with the endoplasmic reticulum, and the inner nuclear membrane^1^. Although the outer and inner nuclear membranes arise from a common membrane, they are functionally distinct membranes. Underlying the inner nuclear membrane is a network of Type V intermediate filament proteins named lamins that provide nuclear rigidity and elasticity^2^. The inner nuclear membrane contains a large number of unique integral inner nuclear membrane proteins^3^, many of which show cell-type-specific expression^4–11^. Inner nuclear membrane proteins function in diverse roles, including nuclear structure, genomic organization, chromatin architecture, gene expression, cell cycle regulation, and cytoskeletal organization^12,1^. The nuclear lamins and its associated inner nuclear membrane proteins define the nuclear lamina.

Emerin is a lamin-binding, integral inner nuclear membrane protein. Mutations in the gene encoding emerin cause X-linked Emery-Dreifuss muscular dystrophy (EDMD1), an inherited disorder causing progressive skeletal muscle wasting, irregular heart rhythms, and contractures of major tendons^13–16^. Evidence suggests the skeletal muscle wasting seen in EDMD is not caused by increased damage to the myofiber, but by impaired differentiation of skeletal muscle stem cells. Supporting this hypothesis, skeletal muscle necrosis and increased skeletal muscle fiber permeability are rarely seen in EDMD patients^17^. Further, emerin knockout mice (also commonly referred to as emerin-null or emerin-deficient mice) exhibit delayed skeletal muscle regeneration and repair, motor coordination defects, and mild atrioventricular conduction defects^18,19^. Skeletal muscle from EDMD1 and EDMD2 patients and emerin-deficient mice both showed altered expression of muscle regeneration pathway components^20,18^. Emerin-deficient myogenic progenitors and emerin-downregulated C2C12 myoblasts exhibit impaired differentiation and myotube formation^21–23^ due to aberrant temporal activation of myogenic differentiation genes^24^ and disruption of key signaling pathways^25^, suggesting defective muscle regeneration contributes to the EDMD skeletal muscle phenotype^22,21,18^.

The coordinated temporal expression of *MyoD, Myf5, Pax3* and *Pax7,* which are important for proper differentiation, was disrupted in emerin-deficient myogenic progenitors^26^ due to the inability of the genome to properly reorganize during differentiation^20,18,25^. This supports the hypothesis that emerin-deficient myogenic progenitors fail to undergo the transcriptional reprogramming required for myogenic differentiation. Furthermore, emerin was shown to bind directly to Histone Deacetylase 3 (HDAC3) and activate its deacetylase activity^27^. HDAC3 activity is required for proper dynamic reorganization of *MyoD, Myf5, Pax3* and *Pax7*^26^. Thus, regulation of HDAC3 activity by emerin is critical for transcriptional reprogramming during myogenic differentiation.

We used histone acetyltransferase (HAT) inhibitors targeting HATs mediating H4K5 acetylation (e.g., Tip60/KAT5) to further test the hypothesis that acetylation dynamics on lysine 5 of Histone 4 (H4K5) were important for in myogenic differentiation. Here we show increased H4K5 acetylation (H4K5ac) contributes to the impaired differentiation of emerin-deficient myogenic progenitors. Targeting H4K5ac dynamics represents a potential new strategy for ameliorating the skeletal muscle wasting seen in EDMD1.

## METHODS

### Pharmacological treatments

We previously showed emerin-deficient myogenic progenitors had impaired differentiation and was rescued by activation of HDAC3^23^. To independently test if altered H4K5 acetylation dynamics was responsible for the impaired differentiation of emerin-deficient progenitors we chose to inhibit HATs. HAT inhibitors (HATi) selected for these studies were chosen because they preferentially inhibit acetylation of lysine residues targeted by HDAC3 (e.g., H4K5)^34^. Cell cycle withdrawal, myosin heavy chain (MyHC) expression and myotube formation were analyzed 36 hours post-differentiation induction, as previously described^23^. HAT inhibitor Nu9056 was selected because it is a highly specific inhibitor of histone acetyltransferase Tip60/KAT5^28^. Tip60/KAT5 mediates the acetylation of H4K5, H4K8, H4K12 and H4K16ac (Table 1). A second HAT inhibitor, L002, was used to test whether inhibition of H4K5 acetylation rescued myogenic differentiation of emerin-deficient progenitors. L002 inhibits H4 acetylation in cells at low micromolar concentrations (Table 1)^29^. A Sirtuin 1 (SIRT1) activator (SRT1720) was used to confirm the rescue of emerin-deficient progenitor differentiation was due to changes in acetylation states of HDAC3 target residues (e.g., H4K5ac). Unlike HDAC3, SIRT1 is an NAD^+^-dependent protein deacetylase^35^. SIRT1 deacetylates H3K9ac, but does not affect H4K5, H4K8 or H4K12 acetylation^36^. 1.5 μM SRT1720 was added to wildtype or emerin-deficient myogenic progenitors upon differentiation induction and differentiation was analyzed after 36 hours (Figure 1A).

**Figure 1.**
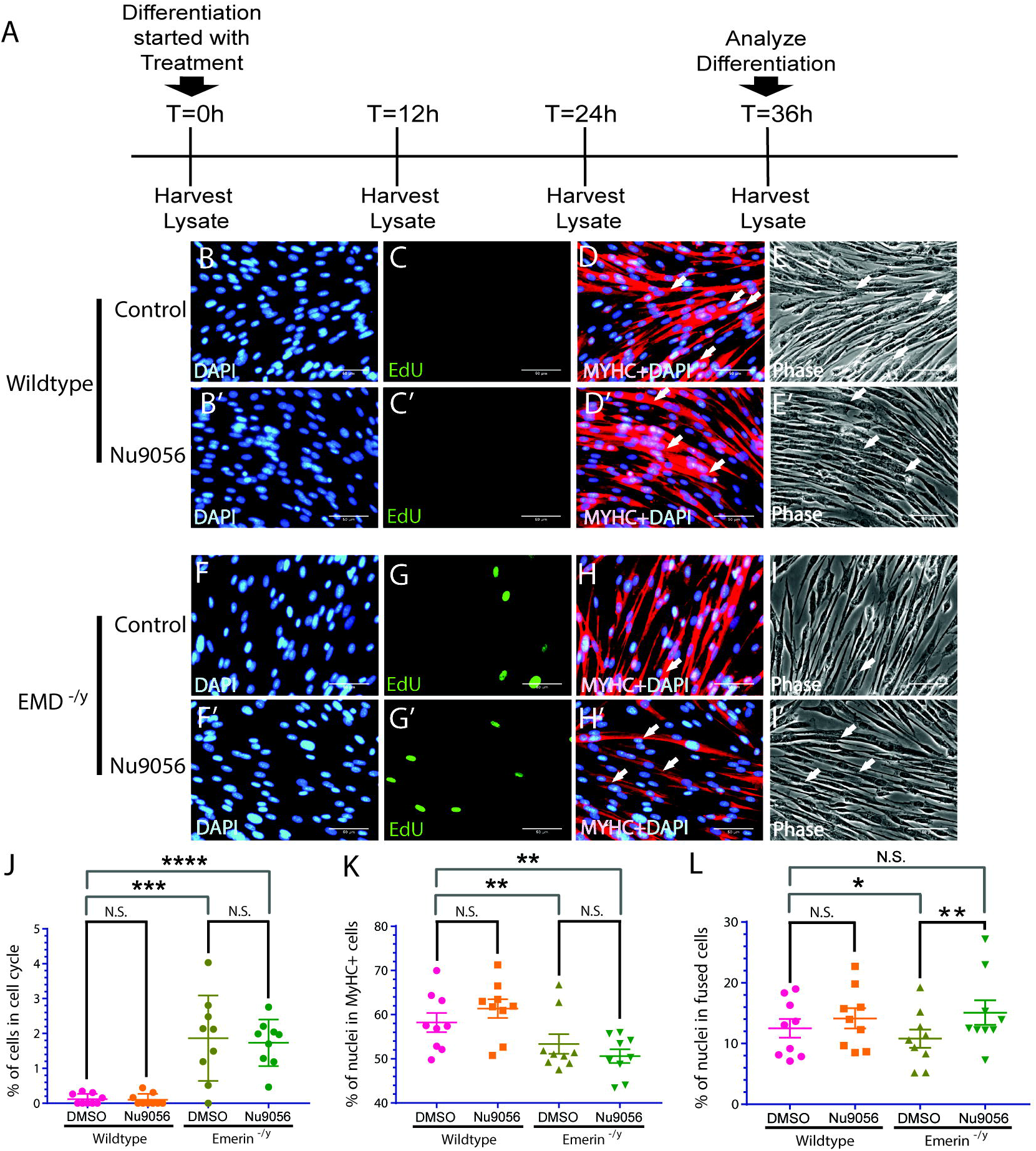
Inhibition of HAT activity with Nu9056 treatment rescues myotube formation in emerin-deficient myogenic progenitors. (A) Timeline showing the time point Nu9056 was added and whole cell lysate collection for western blot analysis. Representative images at 40X magnification of vehicle-treated wildtype (B-E) or emerin-deficient (F-I) and Nu9056-treated wildtype (B’-E’) or emerin-deficient (F’-I’) cells 36 h after initiating differentiation. Arrows mark myotubes (e.g., 3 myotubes in I’ vs 1 myotube in I). (J-L) Quantification of >500 nuclei for each experimental treatment (n=3) was carried out to determine the percentage of myogenic progenitors in the cell cycle (J), expressing MyHC (K) and formed tubes (L) 36 h after inducing differentiation. Results are mean ± s.d. of n=3; N.S., not significant; *, P<0.05; **, P<0.01; ***, P<0.001; ****, P<0.0001 using paired, two-tailed t-tests. Scale bars are 50 μm.

Concentrations of compounds used in this study were based upon the reported half-maximal inhibitory concentration (IC50) for each compound^28–31^ and per manufacturer (EMD Millipore) instructions. 0.25 μM to 30 μM or each compound was added to wildtype and emerin-deficient myogenic progenitors and tested for viability and impaired proliferation. The concentrations used in this study failed to inhibit cell proliferation and had no effect on cell viability. A 1.0 mM stock solution of L002 in DMSO was added to a final concentration of 0.5 μM in differentiation medium. A 1.0 mM stock solution of Nu9056 in DMSO was added to a final concentration of 0.5 μM in differentiation medium. A 3.0 mM stock solution of SRT1720 in DMSO was added to a final concentration of 1.5 μM in differentiation medium. Differentiation media containing each inhibitor or DMSO was added to induce differentiation of wildtype or emerin-deficient myogenic progenitors.

### Cell culture

Wildtype and Emerin-null H2K mouse myogenic progenitors were a generous gift from Tatiana Cohen and Terence Partridge (Children’s National Medical Center, Washington, DC). Emerin-null H2K mice were generated by Tatiana Cohen and Terence Partridge by breeding emerin-null (C57Bl/6) and H-2K^b^tsA58 mice to create emerin-deficient mice in the H-2K^b^tsA58 background^32,33^. Myogenic progenitors were isolated and maintained as previously described^24,33,25^. Briefly, extensor digitorum longus (EDL) muscles were isolated and placed into 2 mg/ml collagenase (Sigma, product #C0130) in DMEM (Invitrogen, product #11995-065) for 1–2 hours at 35°C. Individual fibers were then isolated and each fiber was transferred serially through 2–4 petri dishes containing DMEM to select for undamaged fibers. Fibers were placed into matrigel-coated petri dishes containing DMEM, 10% horse serum (Invitrogen product #16050-098), 0.5% chick embryo extract (Accurate Chemical, product #CE6507), 2% L-Glutamine (Invitrogen, product #25030-081) and 1% penicillin/streptomycin (Invitrogen, product #15140-122) for 3–4 hours at 37°C. Myogenic progenitors were isolated from individual fibers by transferring each fiber into one matrigel-coated well of a 24-well plate containing proliferation media consisting of DMEM, 20% Heat inactivated FBS (Invitrogen, product #10082-147), 2% chick embryo extract, 2% L-glutamine, 1% penstrep and 20 ng/ml γ-Interferon (Millipore, product #IF005). The fibers were incubated for 24–48 hours at 33°C, 10% CO_2_. Upon attachment of a single myogenic progenitor to the well, the fiber was removed and the myogenic progenitor was incubated in proliferation media for another 48 hours at 33°C, 10% CO_2_. Approximately 200 cells are expected after 48 hours and these were split and proliferated until enough cells were obtained for our analyses. H2K myogenic progenitors were maintained in proliferation media at 33°C and 10% CO_2_. Cells between passages 4–10 were used for these studies.

Cell culture of proliferation and differentiation of H2Ks were done as previously described^23^. Briefly, for proliferation, wildtype and emerin-deficient H2K myogenic progenitors were seeded onto tissue culture plates (Falcon cat no. 353046 and 3530003) and maintained at 33°C and 10% CO_2_ in proliferation medium (high glucose DMEM supplemented with 20% heat-inactivated fetal bovine serum, 2% L-glutamine, 2% chick embryo extract, 1% penicillin/streptomycin, sodium pyruvate, 20 units/ml γ-interferon, ThermoFisher Scientific). The plates were coated with 0.01% gelatin (Sigma-Aldrich) prior to seeding.

Wildtype and emerin-deficient H2K myogenic progenitors were seeded onto 12 well tissue culture plates coated with 0.01% gelatin (Sigma-Aldrich) for differentiation induction. Cells were seeded at 23,500 cells/cm^2^ in proliferation media for 24h at 33°C and 10% CO_2_. Differentiation was stimulated by replacing the proliferation medium with differentiation medium (high glucose DMEM with sodium pyruvate, 5% horse serum, 2% L-glutamine, ThermoFisher Scientific). The cells were maintained at 37°C and 5% CO_2_ throughout differentiation.

### EdU assays and immunofluorescence microscopy

Cells were treated with 10 μM EdU (ThermoFisher Scientific) in DMSO 2h prior to fixing, while incubating at 37°C and 5% CO_2_. Cells were then fixed with 3.7% formaldehyde for 15 min and washed three times with PBS. Fixed cells were then stored at 4°C with 0.1% sodium azide in PBS. The cells were permeabilized with 0.5% triton X-100 in PBS for 20 minutes, washed twice with 3% BSA in PBS for five minutes per wash and treated with the Click-IT EdU reaction cocktail for 25 minutes. Cells were washed with PBS and blocked for 1 h at room temperature with 3% BSA with 0.1% Triton X-100. Myosin heavy chain (MyHC) antibodies (1:20, Santa Cruz Biotechnologies, H-300 for L002 and Nu506 experiments; 1:50, Santa Cruz Biotechnologies, B-5 for SRT1720 treatments) were added and the cells were incubated at room temperature for 1 h. The cells were washed with PBS three times and treated with Alexa Fluor 594 secondary antibodies (1:200, C10637; A11032, ThermoFisher Scientific) at room temperature for 1 hour, washed with PBS and incubated with DAPI for 5 minutes.

Images were taken using the EVOS-FL imaging system (ThermoFisher LifeSciences) for experiments with L002. The remainder of the images were taken with the EVOS-FL Auto (ThermoFisher LifeSciences). All images were obtained using a long working distance 40x objective. At least three replicates, with each replicate containing three culture wells per group, were done for each drug treatment. Images from five different sections from each well were taken, with each section containing approximately 50-200 cells. The total number of cells analyzed for each experiment ranged between 500-1500.

The cell counter plugin on ImageJ was used to count proliferating cells. The percent of cells still in the cycle was determined by dividing the number of EdU positive nuclei by the total number of nuclei. The DAPI and MyHC images were superimposed to calculate the percentage of cells expressing MyHC. Myotube formation was determined by superimposing the phase contrast image, which allowed for monitoring nuclei within a shared cytoplasm, with DAPI and MyHC images. Myotube formation, or the differentiation index, was determined by counting the number of nuclei in MyHC positive cells that contained three or more nuclei divided by the total number of nuclei in the field.

### Western Blotting

Differentiating H2K cells were resuspended directly in sample buffer and 50,000 to 100,000 cell equivalents were separated by SDS-PAGE and transferred to a nitrocellulose membrane. The membranes were blocked either at room temperature for 2 h or overnight at 4°C in 3% BSA in PBST (PBS with 0.1% Tween 20). Antibodies against H4 (1:50,000; Millipore, 05-858), H4K5ac (1:1,000; Millipore, 07-327), H3K9ac (1:10,000; Abcam, ab4441), H3K18ac (1:1,000; Abcam, ab1191), H3K27ac (1:1,000; Abcam, ab4729), H4K16ac (1:2,000; Abcam, ab109463), and MyHC (1:1,000; Santa Cruz, B-5) were then incubated either at room temperature for 2 h or overnight at 4°C. The membranes were washed three times in PBS and incubated with Goat Anti-Rabbit HRP or Goat Anti-Mouse HRP secondary antibody (1:10,000; ThermoFisher Scientific) in PBST either at room temperature for 2 h or overnight at 4°C. The membranes were treated with ECL chemiluminescence detection reagent (GE healthcare, product # RPN2106V1 and RPN2106V20 and imaged using the Bio-Rad Chemidoc system (Bio-Rad Laboratories). Densitometry was done using ImageLab software (Bio-Rad Laboratories) as per the manufacturer’s instructions.

## RESULTS

Wildtype and emerin-deficient myogenic progenitors were differentiated for 36 hours in the presence of the HAT inhibitor Nu9056 to independently confirm HAT inhibition rescued emerin-deficient myogenic differentiation. 0.5 μM Nu9056 in DMSO or DMSO alone were incubated with wildtype or emerin-deficient myogenic progenitors upon differentiation induction (Figure 1A). Nu9056 treatment had no effect on cell cycle withdrawal of wildtype or emerin-deficient myogenic progenitors, (Figure 1C, G, J). Nu9056 treatment failed to rescue myoblast commitment, as the number of MyHC-expressing cells was similar in Nu9056-treated (51.0%) and untreated emerin-deficient myogenic progenitors (50.3%; Figure 1H, K). Myotube formation in emerin-deficient progenitors was rescued by Nu9056 treatment, as 15.1% of Nu9056-treated emerin-deficient progenitors fused to form myotubes, compared to 10.8% of DMSO-treated emerin-deficient progenitors (Figure 1I, L). Myotube formation in Nu9056-treated emerin-deficient progenitors was statistically similar to wildtype progenitors (p=0.11).

A second HAT inhibitor, L002, was used to test whether inhibition of H4K5 acetylation rescued myogenic differentiation of emerin-deficient progenitors. Wildtype and emerin-deficient myogenic progenitors were treated with 0.5 μM L002 upon differentiation induction (Figure 1A). L002-treated wildtype progenitors exited the cell cycle normally (Figure 2B’, I). 2.7% of emerin-deficient progenitors failed to exit the cell cycle after 36 hours, as expected (Figure 2F, I). Emerin-deficient progenitors treated with L002 showed a trend toward reducing the number of emerin-deficient cells in the cell cycle (2.1%; p=0.06; Figure 2F’, I). L002 treatment significantly increased the percentage of differentiating emerin-deficient progenitors expressing MyHC (46%, Figure 2G, J; p=0.015). The number of MyHC-positive cells in L002-treated differentiating emerin-deficient progenitors is statistically similar to untreated wildtype progenitors (47.8% in wildtype, p=0.35; Figure 2C, G’, J), indicating rescue of myoblast commitment. L002 treatment increased myotube formation 1.8-fold in differentiating emerin-deficient progenitors (Figure 2H, K) completely rescuing myotube formation to wildtype levels (p=0.97 for L002-treated emerin-deficient cells vs. wildtype cells; Figure 2D, H’, K).

**Figure 2.**
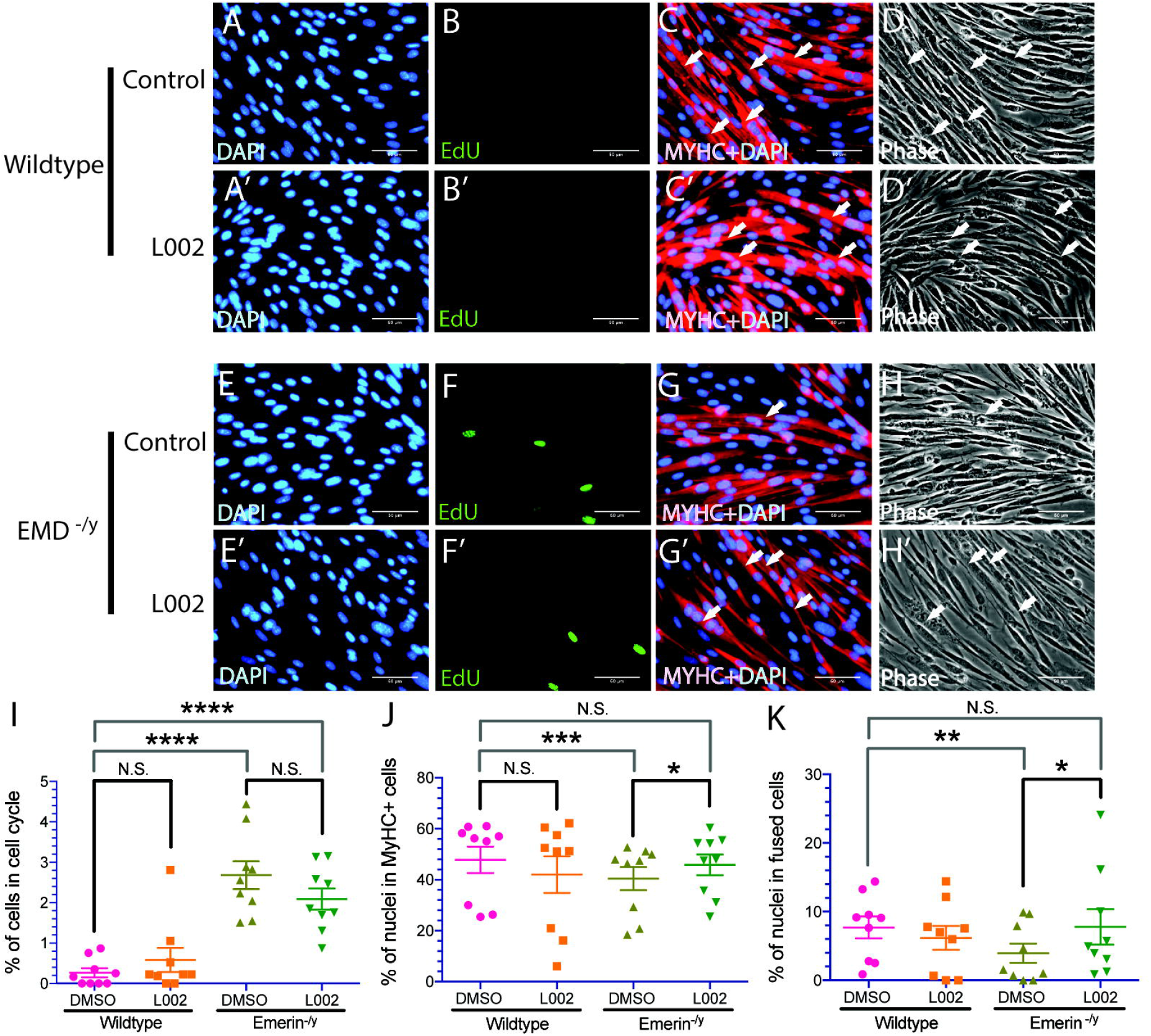
Inhibition of HAT activity with L002 treatment rescues myotube formation and myosin heavy chain expression in emerin-deficient myogenic progenitors. (A-D) or emerin-deficient (E-H) and L002-treated wildtype (A’-D’) or emerin-deficient (E’-H’) cells 36 h after initiating differentiation. Arrows mark myotubes (e.g., 4 myotubes in H’ vs 1 myotube in H). (I-K) Quantification of >500 nuclei for each experimental treatment (n=4) was carried out to determine the percentage of myogenic progenitors in the cell cycle (I), expressing MyHC (J) and formed tubes (K) 36 h after inducing differentiation. Results are mean ± s.d. of n=4; N.S., not significant; *, P<0.05; **, P<0.01; ***, P<0.001; ****, P<0.0001 using paired, two-tailed t-tests. Scale bars are 50 μm.

A Sirtuin 1 (SIRT1) activator (SRT1720) was used to confirm the rescue of emerin-deficient progenitor differentiation was due to changes in acetylation states of HDAC3 target residues (e.g., H4K5ac). Activation of SIRT1 by treatment with 1.5 μM SRT1720 failed to rescue cell cycle withdrawal of differentiating emerin-deficient progenitors, as 7.0% of DMSO-treated and 5.4% of SRT1720-treated cells were cycling (Figure 3F, I; p=0.09). 41.1% of SRT1720-treated differentiating emerin-deficient progenitors expressed MyHC compared to 42.7% of DMSO-treated emerin-deficient progenitors (Figure 3G, J; p=0.49). SRT1720 treatment also failed to rescue myotube formation in emerin-deficient progenitors (Figure 3H, K; p=0.44).

**Figure 3.**
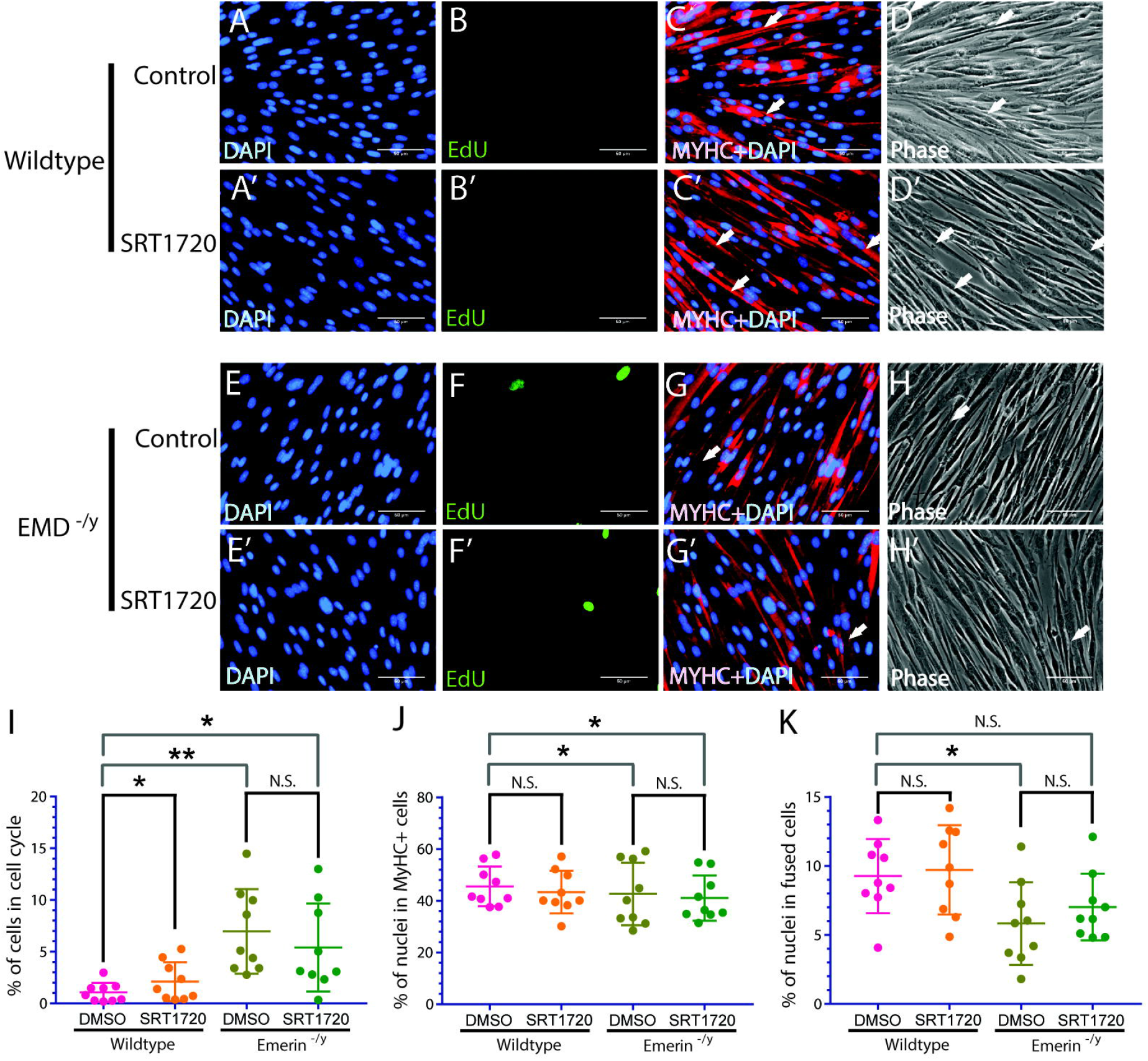
SIRT1 activation with SRT1720 treatment does not change cell cycle exit, myosin heavy chain expression, or myotube formation in emerindeficient myogenic progenitors. (A-D) or emerin-deficient (E-H) and SRT1720-treated wildtype (A’-D’) or emerin-deficient (E’-H’) cells 36 h after initiating differentiation. Arrows mark myotubes (e.g., 1 myotube in H’ and H). (I-K) Quantification of >500 nuclei for each experimental treatment (n=3) was carried out to determine the percentage of myogenic progenitors in the cell cycle (I), expressing MyHC (J) and formed tubes (K) 36 h after inducing differentiation. Results are mean ±s.d. of n=3; N.S., not significant; *, P<0.05 using paired; **, P<0.01 using paired, two-tailed t-tests. Scale bars are 50 μm.

Treatment of emerin-deficient progenitors with L002 during differentiation reduced H4K5ac 3.8-fold (Figure 4A, B), comparable to H4K5ac levels in wildtype progenitors. Nu9056 treatment decreased H4K5ac 3.2-fold in emerin-deficient myogenic progenitors (Figure 4A, B). Decreased H4K5ac seen in emerin-deficient myogenic progenitors treated with Nu9056 is similar to the H4K5ac levels seen in differentiating wildtype progenitors. Western blotting confirmed the levels of H4K5ac were unchanged by treatment with SRT1720 (Figure 4C, D).

**Figure 4.**
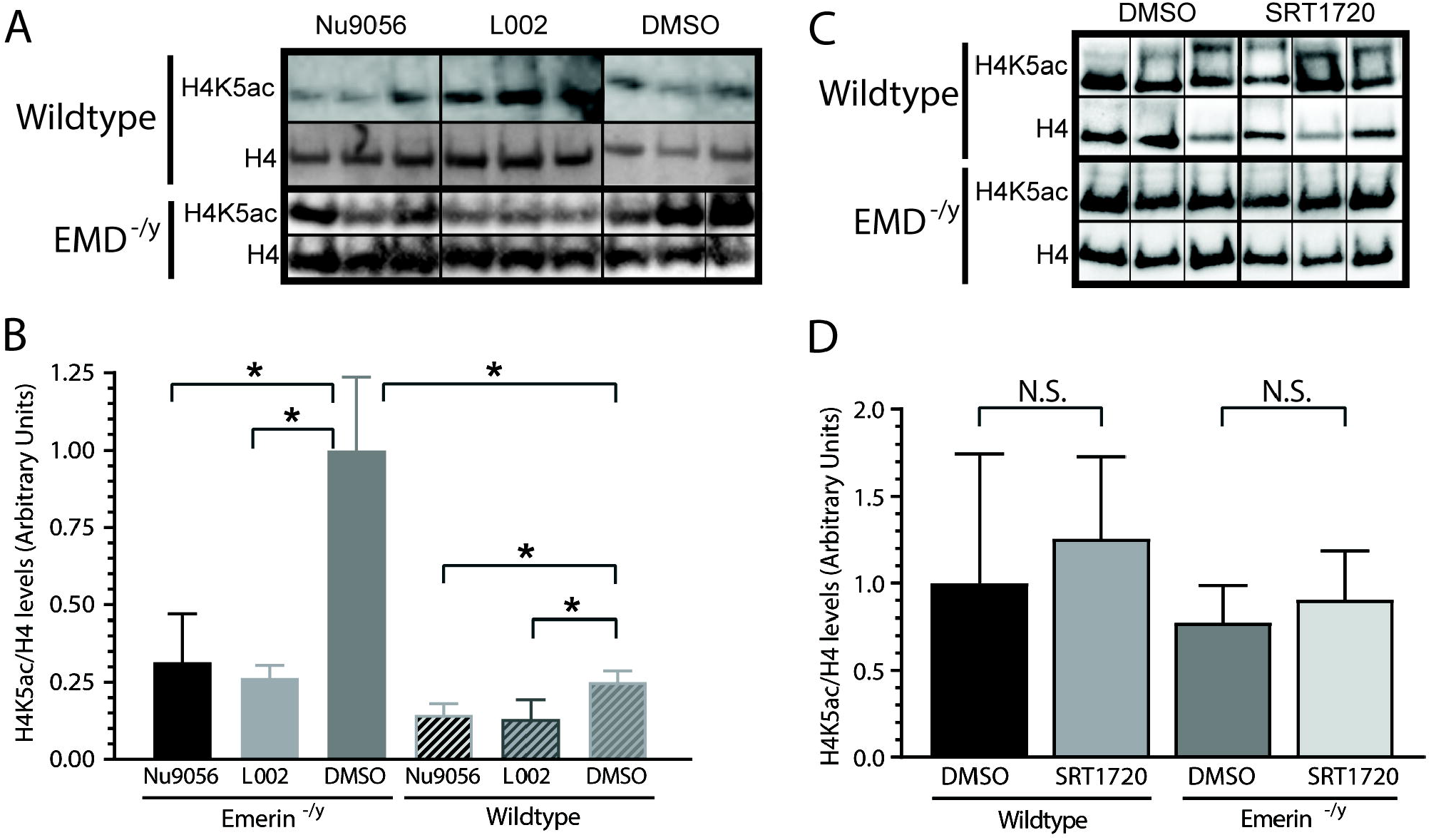
H4K5 acetylation is decreased by treatment with L002 and Nu9056. Western blotting of whole cell lysates treated with (A) Nu9056, L002, or (B) SRT1720 to analyze H4K5 acetylation during differentiation of wildtype and emerin-deficient progenitors. DMSO-only treatment was the control. Three biological replicates are shown for each treatment. (B,D) Densitometry was performed and acetylated H4K5 in each sample was normalized to total H4 protein in each sample. Levels of acetylated H4K5 for each condition were normalized to DMSO-treated cells. Results are mean ± s.d. of n=3 for each condition; N.S., not significant using paired, two-tailed t-tests.

Acetylation of H3K9, H3K18, H3K27 and H4K16 were monitored during impaired differentiation in emerin deficiency in the presence of Nu9056, L002 and SRT1720. H3K9ac, H3K18ac and H3K27ac were all increased in emerin-deficient myogenic progenitors and during emerin-deficient myogenic differentiation (Figure 5), including during the transition to myoblast commitment. Treatment of emerin-deficient progenitors with Nu9056 had no significant effect on acetylation of H3K9, H3K18, H3K27 or H4K16 (Figure 5). L002 treatment had a small effect on H3K18 and H3K27 acetylation. Treatment with the SIRT 1 activator, SRT1720, reduced H3K9ac activity, as expected, since SIRT1 deacetylates H3K9^35,36^.

**Figure 5.**
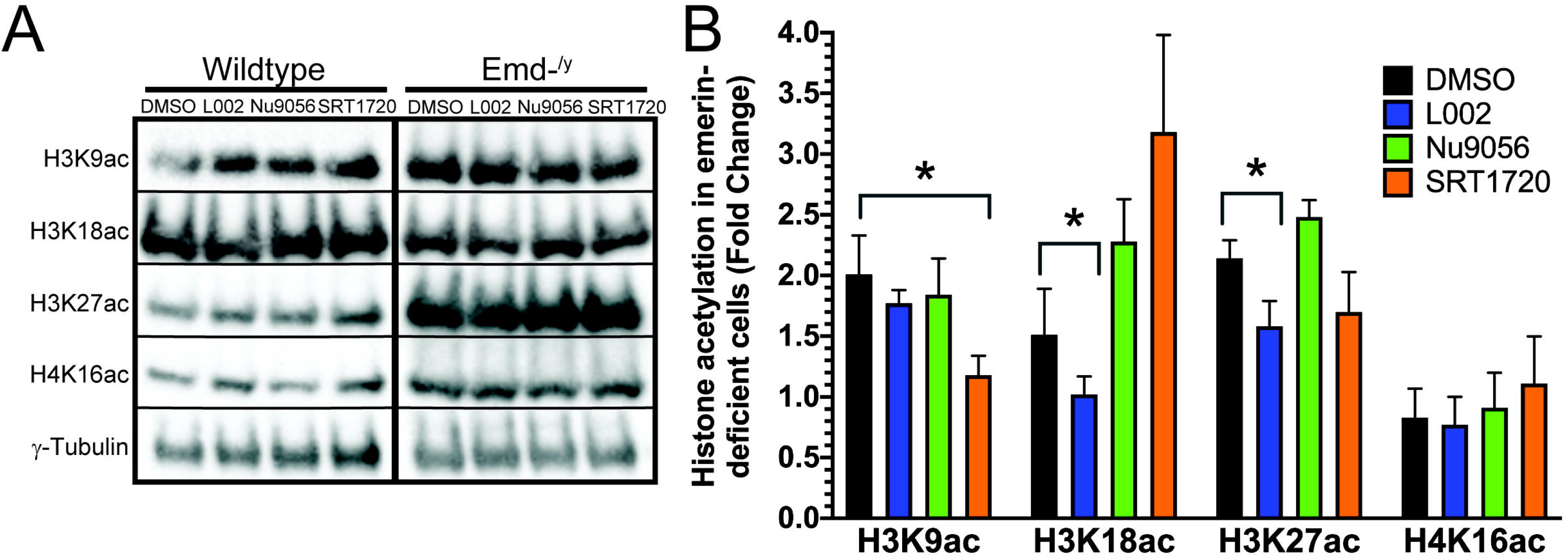
Acetylation of H4K16, H3K9, H3K18 or H3K27 in emerin-deficient myogenic progenitors upon treatment with L002, Nu9056 or SRT1720. Wildtype or emerin-deficient myogenic progenitors were treated with DMSO, L002, Nu9056 or SRT1720 and whole cell lysates were obtained after 36 hours. (A) Western blotting was done with the indicated antibodies to monitor histone acetylation. (B) Quantitation of histone acetylation normalized to γ-tubulin and plotted as fold-change in emerin-deficient cells, as compared to wildtype cells. Results are mean ± s.d. for each condition. N.S., not significant; *P≤0.05 using paired two-tailed t-tests.

## DISCUSSION

The studies presented here used a cell-based system to follow differentiation, in which myotubes are formed by myoblast-to-myoblast fusion or myoblast-to-myotube fusion. Upon stimulation of myogenic progenitors to differentiate, transcriptional reprogramming is initiated, leading to cell cycle exit. This reprogramming activates the myogenic differentiation program and represses the proliferative program, thereby leading to myoblast commitment followed by fusion to form myotubes^37^. Transcriptional reprogramming is compromised in emerin-deficient progenitors^38^. The failure of emerin-deficient progenitors to coordinate the temporal reorganization of their genome during differentiation is predicted to cause this defective transcriptional reprogramming.

Emerin binds directly to HDAC3, the catalytic component of the Nuclear Co-Repressor (NCoR) complex^27,39^ and activates its activity. Emerin-binding recruits HDAC3 to the nuclear envelope. The functional interaction between emerin and HDAC3 coordinates the spatiotemporal nuclear envelope localization of genomic regions containing important transcription factors that control the temporal expression of differentiation genes^27,26^. Loss of emerin disrupts this genomic reorganization resulting in impaired myogenic differentiation. Activation of HDAC3 rescues emerin-deficient myotube formation^26,23^. Nuclear envelope localization of HDAC3 is also important in cardiomyocyte differentiation^40^. In the absence of nuclear envelope-localized HDAC3, repressed genomic loci were aberrantly localized to the nuclear interior resulting in precocious differentiation. Thus, controlling HDAC3 nuclear envelope localization and activation is an important regulatory mechanism used for differentiation.

The results presented here support the role of emerin in controlling histone acetylation dynamics by regulating HDAC3 activity. Using HATi specifically targeting acetylation of residues deacetylated by HDAC3 (e.g., H4K5ac), we found HAT inhibition rescued impaired differentiation in emerin deficiency. This recapitulated the rescue seen by treatment of emerin-deficient progenitors with an HDAC3 activator. Thus, H4K5 acetylation dynamics are predicted to be important for ensuring proper transcriptional reprogramming upon differentiation induction (Figure 6). Similar to HDAC3 activation, HDAC inhibition primarily affected later differentiation transitions^23^, suggesting emerin regulation of HDAC3 activity controls the temporal expression of these later genes. This may result from failure to completely reprogram the transcriptome upon differentiation induction or by specifically regulating the latter steps of the gene expression program.

**Figure 6.**
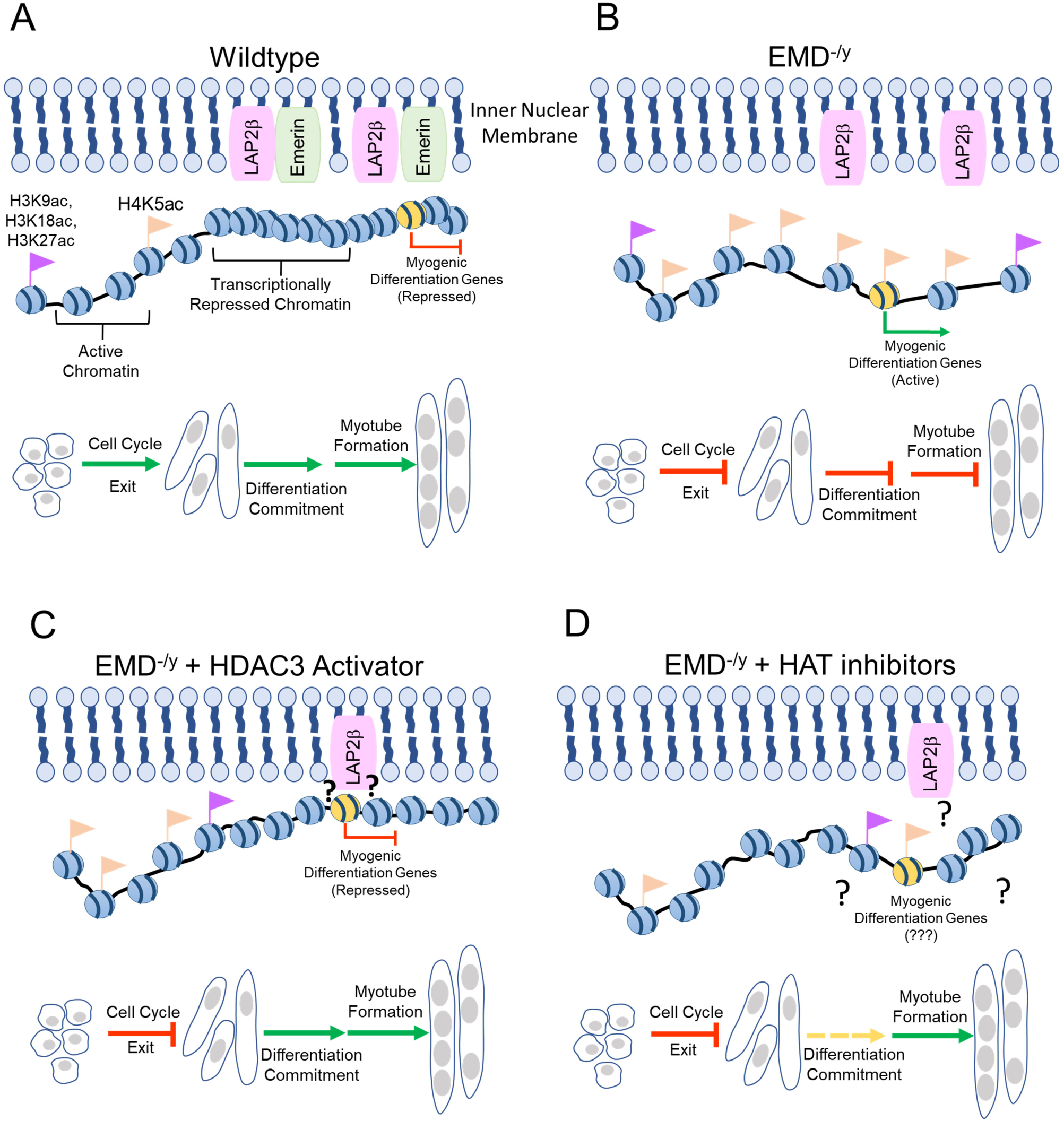
Effects of altered H4K5 acetylation dynamics on myogenic differentiation. (A) Wildtype myogenic differentiation. (B) Lack of emerin results in impaired differentiation with loss of Myf5 localization and increased H4K5 and H3 acetylation states. (C, D) Treatment with HAT inhibitors (C) and HDAC3 activators (D) restore the H4K5 and H4K5ac equilibrium and rescue myotube formation with no effect on cell cycle exit. (C) HAT inhibitor L002 partially rescues myoblast commitment by targeting unknown histone modifications to alter chromatin architecture at the nuclear envelope. (D) Treatment with HDAC3 activators induces Myf5 nuclear envelope localization. Red arrows indicate impaired differentiation programming, solid dark green arrows indicate normal differentiation programming, and dashed light green arrow signifies partially impaired differentiation.

Nu9056 exhibits high specificity, as the half-maximal inhibitory concentration (IC50) of Nu9056 for Tip60/KAT5 is 20-40-fold lower than for the histone acetyltransferase p300 or the histone acetyltransferase pCAF/GCN5 (Table 1). L002 was identified as a p300-specific inhibitor^29^. p300 acetylates H3K18 and H3K27; it has also been reported to acetylate H4K5, H4K8, H4K12 and H4K16^41,42^. L002 was previously reported to decrease H4 acetylation^29^, but this study was the first to show L002 inhibits acetylation of H4K5. Although H3K9ac, H3K18ac and H3K27ac were increased in emerin-deficient myogenic progenitors, treatment with Nu9056 had no significant effect on acetylation of H3K9, H3K18 or H3K27; L002 treatment had a small effect on H3K18 and H3K27 acetylation; SRT1720 only affected H3K9 acetylation (Figure 5). Collectively, the distinct specificities of these compounds at the concentrations used in this study demonstrates that changes in acetylation of H3K9, H3K18, H3K27 and H4K16 cannot be responsible for the rescued myotube formation. Rather, these results suggest rescue by L002 and Nu9056 occurs primarily through rescuing H4K5 acetylation dynamics. This is consistent with our previous studies using HDAC3 inhibitors and activators^23,26^. It is possible L002 may act to rescue impaired myogenic commitment by p300-mediated acetylation of H3K18 or H3K27 (Figure 6). Alternatively, L002 may act through an unknown mechanism or act on other HATs to rescue myoblast commitment, as L002 appears to be more promiscuous at lower concentrations. Targeting these dynamic epigenetic changes represents a new potential strategy for ameliorating the skeletal muscle wasting seen in EDMD1.

It is important to elucidate how emerin regulates the dynamic epigenetic changes occurring during myogenic differentiation to control the transcriptional programs needed for passage through specific transition points. HAT inhibition and HDAC3 activation successfully rescued the latter steps of emerin-deficient myogenic differentiation (this study)^23,26^, suggesting emerin regulation of H4K5ac dynamics during transcriptional reprogramming likely impairs the expression of genes acting at later stages of differentiation (e.g., myotube formation). Consistent with these results, HDAC3 inhibition by RGFP966 blocked MyHC expression and myotube fusion in both differentiating wildtype and emerin-deficient myogenic progenitors^23^. We propose H4K5 acetylation levels are tightly regulated and that increases or decreases in H4K5 acetylation levels impairs the transition from committed, differentiating myoblasts, to myotubes by altering transcription reprogramming upon differentiation induction. Collectively, our results support pharmacological targeting of H4K5 acetylation as a potential therapeutic strategy for rescuing muscle regeneration in EDMD.

## Acknowledgements

We thank the members of the Holaska laboratory for the many helpful discussions regarding these studies and preparation of this manuscript. This study was supported by the National Institute of Arthritis and Musculoskeletal and Skin Diseases of the National Institutes of Health under Award Number R15AR069935 (to J.M.H.). The content is solely the responsibility of the authors and does not necessarily represent the official views of the National Institutes of Health.

## Abbreviations

BSA: Bovine Serum Albumin
DAPI: 4’,6-diamidino-2-phenylindole
DMEM: Dulbecco’s Modified Eagle’s medium
DMSO: Dimethyl Sulfoxide
ECL: Enhanced Chemiluminescence
EDL: Extensor Digitorum Longus
EDMD: Emery-Dreifuss Muscular Dystrophy
EdU: 5-Ethynyl-2’-deoxyuridine
FBS: Fetal Bovine Serum
H3K9: Histone 3 lysine 9
H3K9ac: Histone 3 acetylated on lysine 9
H3K18: Histone 3 lysine 18
H3K18ac: Histone 3 acetylated on lysine 18
H3K27: Histone 3 lysine 27
H3K27ac: Histone 3 acetylated on lysine 27
H4: Histone 4
H4K5: Histone 4 lysine 5
H4K5ac: Histone 4 acetylated on lysine 5
H4K16: Histone 4 lysine 16
H4K16ac: Histone 4 acetylated on lysine 16
HAT: Histone Acetyltransferase
HATi: Histone Acetyltransferase inhibitor
HDAC: Histone Deacetylase
HDACi: Histone Deacetylase Inhibitor
HRP: Horseradish Peroxidase
IC50: Half-maximal Inhibitory Concentration
NCoR: Nuclear Co-Repressor
MyHC: Myosin Heavy Chain
PBS: Phosphate Buffered Saline
PBST: Phosphate Buffered Saline with Tween
SIRT1: Sirtuin 1

