## Supplemental Table 1 for "Histone acetyltransferase inhibition rescues differentiation of emerin-deficient myogenic progenitors"

**Table 1. Histone modifying enzymes and the drugs used to target their activity used in this study.**

| **Enzyme targeted** | **Histone**  **residue**  **acetylated** | **Concentration for half-maximal activity for HAT inhibitors or SIRT activator (μm)** | **Concentration used in this study** |
| --- | --- | --- | --- |
| **Tip60 (HAT)** | H4K5, H4K8, H4K12, H4K16 | Nu9056 (<2.0 μm) | 0.5 µm |
| **p300 (HAT)** | H3K18, H3K27 | L002 (1.98 μm) | 0.5 µm |
|  | H4K5, H4K12, H4K8, H4K16, | L002 (0.3 µm; cell-based assay) | 0.5 µm |
|  | H3K9, H3K14, H3K23 | ? | 0.5 µm |
| **pCAF/GCN5 (HAT)** | H3K9 | L002 (34 μm) | 0.5 µm |
| **SIRT1 (HDAC)** | H3K9 | SRT1720 (0.16 μm) | 1.5 µm |
|  | H3K14, H4K16, H1K26 | ? | 1.5 µm |

Abbreviations: HAT, histone acetyltransferase; HDAC, histone deacetylase; H1K26, histone 1, lysine 26; H3K9, histone 3 lysine 9; H3K14, histone 3 lysine 14; H3K18, histone 3 lysine 18; H3K23, histone 3 lysine 23; H3K27, histone 3 lysine 27; H4K5, histone 4 lysine 5; H4K8, histone 4 lysine 8; H4K12, histone 4 lysine 12; H4K16, histone 4 lysine 16.
